# Motor expertise facilitates the precision of state extrapolation in perception

**DOI:** 10.1101/158469

**Authors:** Nicolas Ludolph, Jannis Plöger, Martin A. Giese, Winfried Ilg

## Abstract

Predicting the behavior of objects in the environment is an important requirement to overcome latencies in the sensorimotor system and realize precise actions in rapid situations. Internal forward models that were acquired during motor training might not only be used for efficiently controlling fast motor behavior but also to facilitate extrapolation performance in purely perceptual tasks. In this study, we investigated whether preceding virtual cart-pole balancing training facilitates the ability to extrapolate the pole motion. We compared a group of 10 subjects, proficient in performing the cart-pole balancing task, to 10 naïve subjects. Our results demonstrate that preceding motor training increases the precision of pole movement extrapolation, although extrapolation is not trained explicitly. Additionally, we modelled subjects’ behaviors and show that the difference in extrapolation performance can be explained by individual differences in the accuracy of internal forward models. When subjects are provided with feedback about the true pole movement in a second phase, both groups improve rapidly. The results indicate that the perceptual capability to extrapolate the state of the cart-pole system accurately is implicitly trained during motor learning. We discuss these results in the context of shared representations and action-perception transfer.

## Introduction

Expert tennis players are able to extrapolate the motion of a tennis ball and return it skillfully in order to score a point solely based on their present percept. The current consensus is that internal forward models, which predict the dynamic behavior of the body and objects in the environment (such as the ball and tennis racket) support the control of movements [1–3]. Especially in fast situations internal forward models seem to be exploited to overcome the delay of sensory input [4,5] and to predict events [6–9]. Since motor control is inherently related to the prediction of sensory consequences in order to act optimally, the question arises whether motor expertise facilitates the process of perceptual state extrapolation when asked to explicitly report the state as precise as possible and how motor expertise determines the accuracy of such extrapolations.

In past studies, subjects’ general ability to predict the behavior of objects in order to act purposefully has been investigated using diverse paradigms. For example, La Scaleia et al. [10] have examined subjects’ ability to intercept partially occluded ball trajectories. Despite the occlusion, subjects were able to intercept the ball with high precision. In another paradigm subjects learned to control objects with internal degrees of freedom [11], such as mass-spring-damper objects [12], a virtual cup of coffee [13] or, very recently, the cart-pole system [14]. Predicting the behavior of these objects is non-trivial, because all exhibit a high degree of complexity. Mehta and Schaal [15] have examined subjects while controlling the cart-pole system with the goal to balance the pole. They compared subjects’ actions under full visual feedback to actions during short occlusions of up to 550 milliseconds. Despite missing visual feedback, subjects’ actions were indistinguishable from those under full vision. In correspondence with the above-mentioned studies they concluded, that actions during the occlusion are performed based on an extrapolated state, which replaces the missing visual feedback. In line with this conclusion, we showed that subjects learn to perform their actions in the cart-pole balancing task predictively [14], suggesting that subject implicitly extrapolate the systems behavior to plan and time the actions in advance. Moreover, if the dynamics of the system were changed, subjects needed to adapt the action timing to the changes just as it is known from sensorimotor adaptation paradigms [16]. This suggests that an internal forward model, which mimics the cart-pole dynamics, is adapted and, furthermore, that it is used to extrapolate the state during occlusion of the pole in Metha and Schaal’s experiment [15]. Altogether, these studies indicate that motor training alters not only motor control but also affects mechanisms of sensory processing. However, these studies only provide evidence in favor of improved state extrapolation capabilities for controlling motor behavior.

Psychophysical studies suggest that internal simulations are also exploited during different tasks of perceptual motion extrapolation. Graf et al. [17], for example, showed subjects human actions as point-light movies and asked them to judge whether a static posture, shown after an occlusion of varying duration, is a plausible succession of the action. The static postures either matched with the duration of the occlusion or not. They found, that subjects’ responses match with the internal simulation hypothesis, which predicts that subjects’ error rates depend on the mismatch of the shown posture and the duration of the occlusion. Since we are all acquainted with human motion, comparing different levels of motor expertise was not the goal of the study but the study shows that humans can be very precise in spatial-temporal extrapolation. Aglioti et al. [18], in comparison, examined how the expertise in playing basketball influences the ability to judge the success of a free shot. They found that elite basketball players are more accurate in this task than subjects with similar visual but with much less motor experience (coaches and sports journalists). While these studies show that internal simulation and motor expertise influence non-motor related discriminative decisions, it is still an open question whether motor expertise improves the precision in extrapolating the state of the controlled system. Predicting the exact state of a complex dynamical system, such as the cart-pole system, over a certain duration of visual occlusion and reporting it explicitly, is presumably a considerably more challenging task than, for example, only judging the success of a free shot.

In this study, we examined whether prior motor experience facilitates the accuracy in predicting the state of the cart-pole system not only implicitly for the purpose of control but also explicitly for reporting the state. We hypothesized that subjects who are able to balance the cart-pole system are more accurate in extrapolating the pole angle dynamics than subjects without motor expertise who are only visually familiar with the cart-pole system. Furthermore, we hypothesized that the reason for the enhanced ability to predict the pole angle dynamics precisely is that motor experienced subjects possess a more accurate internal model of the cart-pole system. Specifically, we hypothesized that the time horizon, over which accurate predictions can be performed, is larger for subjects with motor expertise and fitted subjects’ responses with a corresponding model. Lastly, we show that subjects without motor expertise are able to improve in extrapolation performance when provided with feedback. Our results are in line with the hypothesis, that motor training facilitates perceptual capabilities and further suggest that these perceptual capabilities are not only implicitly accessible for the purpose of control but also to perform explicit and precise state extrapolation when asked to report the state in a perceptual task.

## Methods

### Subjects

Twenty healthy young participants (age range 18-33 years, mean age 26.1) participated in the main experiment (**Figure 1**). All participants gave informed written consent prior to participation. The study had been approved by the local institutional ethical review board in Tübingen (AZ 409/2014BO2). Of the twenty participants, ten subjects participated in previous experiments in our laboratory and were thereby able to control the cart-pole system (skill acquisition). Consequently, these subjects were assigned to the motor control familiar (MF) group. The remaining ten subjects (group VF) were naïve regarding the control of the cart-pole system. All participants were naïve regarding this study. For the motor control familiar subjects the average time between the skill acquisition and the present experiment was 335 days. Gender and age have been balanced between groups (mean ±sd: MF 26.3 ±3.2 years, VF 25.9 ±3.5 years). All participants had normal or corrected to normal vision. Subjects were paid 8 Euros per hour independent of their performance.

### Experimental Protocol

We examined two groups representing different degrees of motor expertise (experienced vs. unexperienced). In the first group (motor familiar, MF), subjects had already learned to balance the pole on the cart in a previous study and were therefore familiar with the cart-pole system. Subjects in the second group (visual familiar, VF) had no prior exposure to the cart-pole system and were visually familiarized in dedicated blocks during this study. Subjects of both groups performed overall 11 blocks of the cart-pole extrapolation task (**Figure 1**A). Feedback about the real pole position was only provided in the last five blocks. The very first block (block E) was used to ensure task understanding. The baseline prediction performance was assessed during the second block (block B). Each of the subsequent four blocks (T1-T4) was preceded by a 5 minutes long motor control or visual familiarization block (**Figure 1** A) depending on the group affiliation. Subjects in the group MF only performed motor control blocks while subjects in the group VF only performed visual familiarization blocks. Subsequent to the familiarization phase (T1-T4), participants in both groups performed the extrapolation task for another five blocks (F1-F5), in which also feedback was provided after every response (feedback phase, (**Figure 1**B). Thus, the prediction accuracy could be improved by the means of error correction. The whole experiment lasted about 90 minutes.

**Figure 1.**
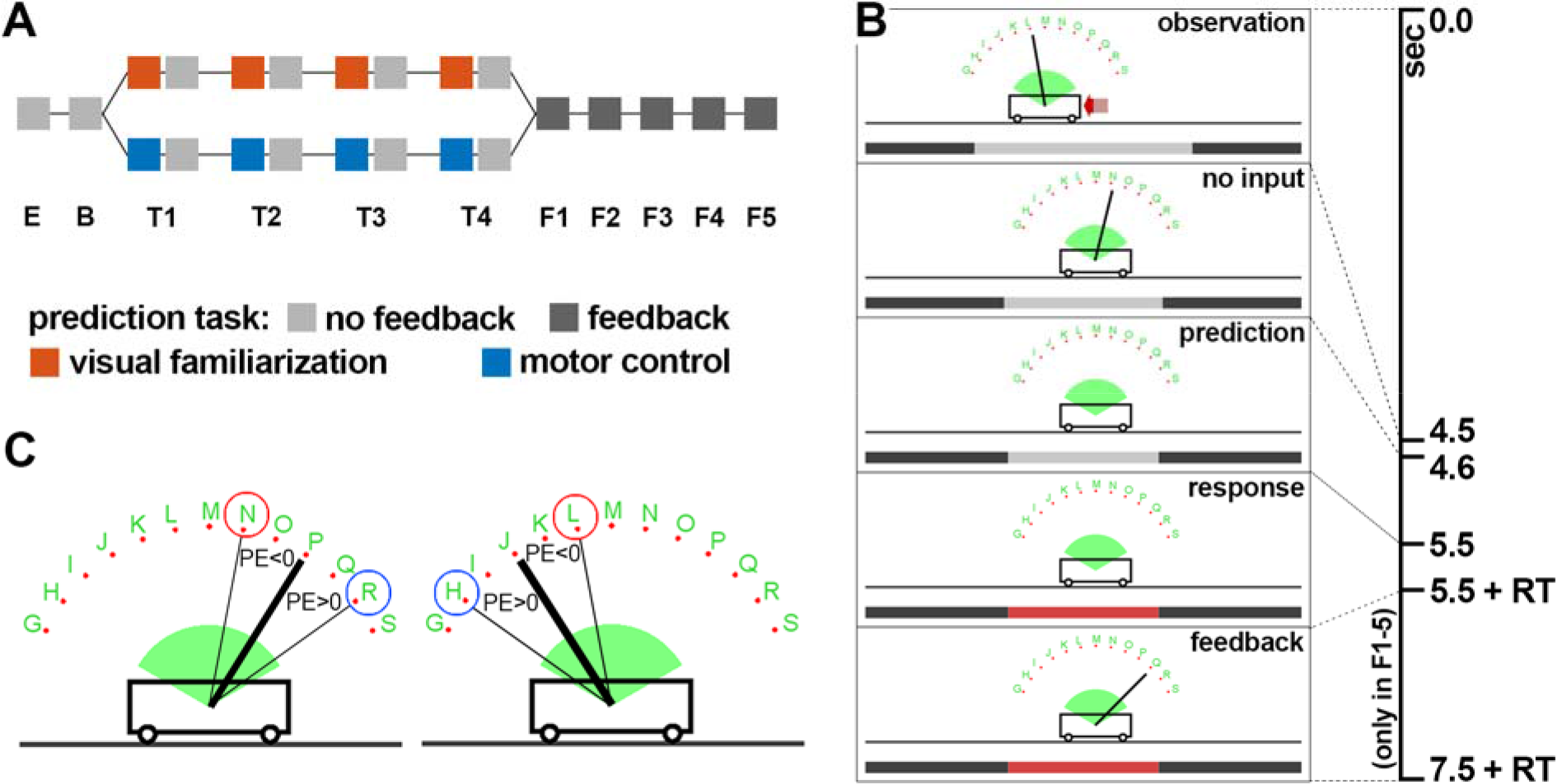
Cart-pole extrapolation task. (A) Phases of each trial during the cart-pole extrapolation task. During the observation phase, a red arrow indicated the direction and magnitude of the force, which was applied during balancing. In the prediction phase, even though the pole was hidden, the dynamical system was simulated further. Thus, the cart and pole kept moving. Subsequently the time bar turned red and the subject gave a response (keypress) representing the expected location of the pole. The feedback phase indicated the correct pole angle and was optional (see below). RT: response time. (B) Sequence of the blocks and tasks. Subjects in the group VF (and extVF) followed the upper (orange), while subject in the group MF followed the lower (blue) path. In block E, the extrapolation task is explained, while in block B, the baseline performance is measured. The subsequent four blocks are preceded by a visual familiarization block (orange) or motor control block (blue). Feedback was provided during the last five blocks (dark gray). (C) Schematic illustration of the prediction error regarding the pole angle. Notice that an underestimation of the pole movement (blue) corresponds to a positive error irrespective of the side.

### Experimental Setup

Visual feedback was provided on a 17 inch monitor (1920×1200px) using the Psychtoolbox [19–21] for MATLAB (The Mathworks, Inc.) at a refresh rate of 60 Hz (**Figure 1** and **Figure 2**). Subjects’ heads were supported using a chin-rest 60 cm away from the screen. The pole of the virtual cart-pole system was rendered as a 160 pixels long line, corresponding to about 3.01cm on the screen or 2.92° in visual space. A keyboard for recording subjects’ responses during the prediction and visual familiarization blocks was positioned between the subject and the monitor. For subjects in the group MF, a SpaceMouse^®^ Pro (3Dconnexion) was additionally placed next to the keyboard for controlling the cart-pole system in the motor control blocks. The cart-pole dynamics [22] are described by the pole mass (0.08 kg), pole length (1 m), cart mass (0.4 kg) and gravitational constant (3.5 m/s^2^). We did not simulate friction. Like in our previous study [14], we implemented the simulation in MATLAB (The MathWorks, Inc.) using the 4^th^-order Runge-Kutta method.

**Figure 2.**
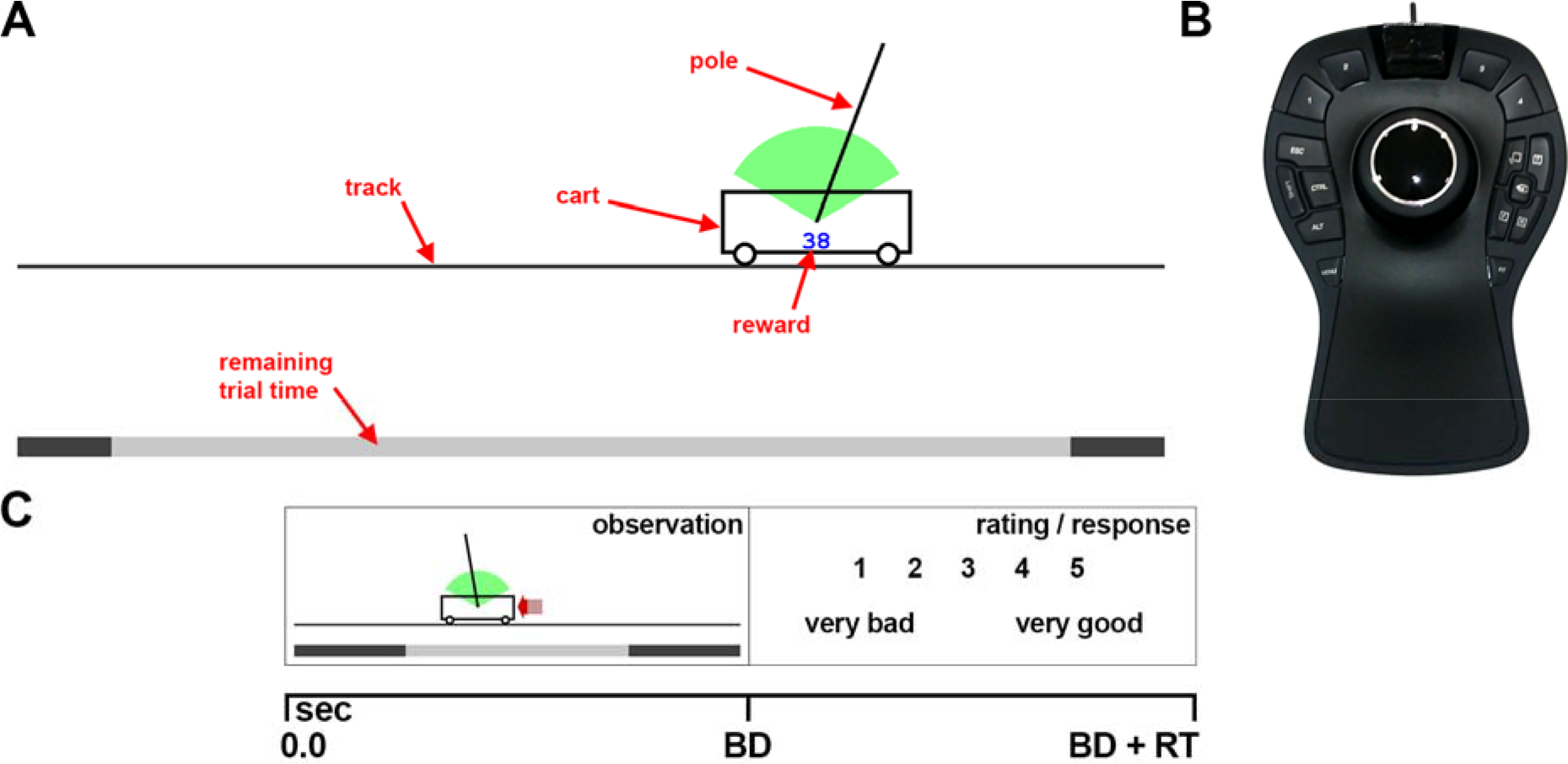
Motor control and visual familiarization task. (A) Visualization of the cart-pole system for both tasks. In contrast to the motor control blocks, in the visual familiarization blocks the force was indicated as red arrow (like in the extrapolation task). (B) The input device, which was used for controlling the system during the motor control blocks. The knob of the input device can be shifted left and right, which was used to control the virtual force that is applied to the cart from either side. (C) Phases of each trial during the visual familiarization. During the observation phase, a balancing attempt was shown that lasted up to 30 seconds. Afterwards, subjects rated the attempt on a scale from one (very bad) to five (very good). BD: balancing duration, RT: response time.

### Cart-Pole Extrapolation Task

In the extrapolation task, participants were asked to indicate the expected angle of the pole after a short occlusion (900ms) during the simulation of the cart-pole system (**Figure 1** B). Every trial began with a short sequence (4.5sec) extracted from previous balancing attempts of other subjects (**Figure 1** B). Afterwards the system was simulated over one second with the input force set to zero. Within this second, the pole was occluded after 100ms for the remaining time of 900ms. Since the input force is zero during this time, there are neither any discontinuities nor any external sources influencing the systems behavior even 100ms before the occlusion, making deterministic extrapolation of the pole dynamics without additional information possible. To indicate the expected pole angle, subjects had to choose one of thirteen response options (**Figure 1**B, C), which were evenly distributed over the pole angle range [-65°, 65°] and identified by nearby letters. Each letter corresponds to the key subjects had to press for choosing the respective response option. Subjects were instructed to always choose the response option that is closest to the expected pole angle. A time bar (**Figure 1**B) indicated the time progression within the trial. Subsequent to observing the system’s movement for overall 5.5 seconds, the time bar turned red which was the sign for the subjects to respond. Depending on the block (**Figure 1**A), subjects received feedback after having responded, which consisted of the presentation of the true pole position. We presented in every block the same 40 balancing attempts (stimuli) in pseudorandom order and recorded subjects’ responses. The stimuli were selected such that correct responses were balanced between the left and right side, while additionally covering a variety of pole angles and angular velocities at the time of occlusion onset.

### Familiarization Tasks

During the motor control blocks, participants had to balance the cart-pole system (**Figure 2**A, for details see also [14]). Specifically, they were asked to balance the system for a maximum of 30 seconds without letting the pole fall out of the green arc and without driving off the track. They therefore had to apply lateral virtual forces to the cart, which in turn accelerated the cart. In order to control the force, they used an input device (SpaceMouse^®^ Pro, 3Dconnexion) with a lateral degree of freedom in displacement (see Experimental Setup and **Figure 2**B). Pushing the knob of this device to the side was translated into a proportional force into the same direction. Because the device position was aligned with the monitor, the left-right knob movement was in correspondence with the virtual force and cart movement on the monitor. Hence, a rightward knob movement caused a force that pushed the cart to the right. Subjects quickly remembered how to balance the pole on the cart and therefore rapidly reached high performance (see Supplementary Material).

In the visual familiarization blocks (**Figure 2** C), participants had to observe balancing attempts of previously recorded subjects instead of controlling the system themselves. Like in the extrapolation task, a red arrow indicated the force, which had been applied during actual balancing. Apart from these arrows, visual information was identical in both conditions (see Supplementary Material). In order to ensure that participants pay attention to the balancing performance and to the dynamic behavior of the cart-pole system, they had to rate every balancing attempt on a scale from one (very bad) to five (very good). Participants were instructed that the task of the individuals, whose balancing performance is shown, was to balance the pole on the cart for a maximum of 30 seconds and, that the persons were rewarded (points) depending on the duration and proficiency of the balancing attempt. Importantly, the presented attempts during these blocks were different from those shown during the extrapolation task. Additionally, instead of only showing short sequences as in the extrapolation task, the entire balancing attempts were shown, which each lasted up to 30 seconds.

## Control Experiment

In order to examine whether subjects improve in prediction accuracy if the visual familiarization phase exhibits a similar overall duration as the preceding motor control training of the other group, we performed a small control experiment (group extVF). In this condition, each visual familiarization block was 20 minutes long (instead of 5 minutes), providing the participant with an overall interaction time of 80 minutes with the system. This is comparable to the duration which subjects in the group MF needed during the initial motor training to master the cart-pole balancing task. It turned out that this procedure was quite cumbersome and annoying for the subjects, not only because of the overall duration but also because subjects did not improve without receiving feedback (ANOVA which examined the effect of the blocks B and T1-T4 on the prediction error, within-subject factor block: p=0.92). Hence, we only recorded three subjects in control experiment. Although a statistical comparison between groups is not possible, subjects in this group exhibited a similar average prediction error (blocks B and T1-T4) as subjects in the group VF (mean ±sd: extVF −14.84° ±1.92, VF −14.84° ±2.94). We therefore concluded for the further analysis that there is no difference between repetitively observing and rating balancing attempts in blocks of 5 or 20 minutes duration.

## Data Analysis

Data preprocessing has be performed in MATLAB (The Mathworks, Inc.). Due to a recording error, one of the subjects in the group MF had to be excluded, leaving overall 9 subjects in the group MF and 10 in the group VF. All statistical analyses have been performed in R (v3.3.2) using the packages lme4 (v1.1), lmerTest (v2.0), phia (v0.2) and nnet (v7.3). Measures have been examined regarding normality using visual inspection of quartile-quartile plots.

### Rating of balancing attempts during visual familiarization

Based on participants’ responses during the visual familiarization, we investigated how the two parameters balancing duration (BD) and mean absolute pole angle (maPA) influenced the rating. While the balancing duration is an obvious measure of balancing performance, the mean absolute pole angle represents how easeful the balancing was performed. The lower the mean absolute pole angle, the smoother the system was controlled and the more vertical the pole was held throughout the trial. First, we examined the effect of the rating on the two parameters using a mixed-effects ANOVA. This provides information about the difference between the rating classes in these parameters. Secondly, we fitted a multinomial log-linear model that mapped the two parameters (BD, maPA) to the rating. Using model comparison (likelihood ratio test), we determined the significance of each of the two parameters (and their interaction) for rating. Thereby, we revealed that both parameters and their interaction are influencing the rating of the balancing attempts significantly (see Results).

### Pole angle prediction error

The main measure for the accuracy in predicting the pole angle is the prediction error (**Figure 1**C). In the extrapolation task, subjects chose in every trial one of 13 response options corresponding to the location where they expected the pole to be after the occlusion. We defined the prediction error (PE) as the angular difference between the true pole position and subject’s response multiplied by the sign of the true pole angle (Equation 1,(**Figure 1**C).

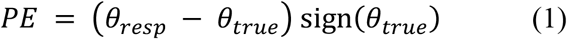

Thus, negative errors correspond on both sides to an underestimation of the pole’s downwards motion, while positive errors correspond to an overestimation. Statistical examination of the factors influencing the prediction error was performed using linear mixed-effect ANOVAs. In all models, we accounted for individual performance and learning rates by introducing random effects per subject.

We first examined whether the ability to extrapolate the pole angle improves during the familiarization phase (B, T1-T4). Since we did not find any improvement (see Results), we then compared the prediction error between groups in the blocks T4 (no feedback, but familiar) and F5 (feedback). These blocks represent different knowledge levels: (T4) subjects are familiar with the system, but did not receive feedback yet, and (F5) subjects had time to utilize the feedback for improvement. We conducted a mixed-effect ANOVA that examined the effect of group and feedback on the prediction error in these blocks.

### Models of inaccurate state predictions

In attempt to identify the cause for subjects’ inability to predict the pole angle precisely, we investigated whether an inaccuracy of the subjects’ individual internal forward models could account for the difference in observed prediction errors. Specifically, if subjects had a perfect forward model (PERF) of the pole dynamics, the prediction error would be roughly zero (neglecting the discretization error introduced by the finite number of response options). Simplification of the dynamics, such as assuming constant pole acceleration (cACC, Supplementary Material) or constant pole velocity (cVEL, Supplementary Material), as it might be appropriate for free-falling objects, introduces a considerable error in the extrapolated pole angle because the actual pole acceleration depends on the angle. Subjects may also be able to predict the pole movement precisely only over the first few milliseconds, before they have to switch to a heuristics because of their limited computational capabilities. Our hypothesis is that motor familiarity facilitates these capabilities, leading to an increased duration of the interval for which precise extrapolation is possible (extrapolation horizon). We modelled this behavior by a class of partly heuristic response models, which are described in more detail in the Supplementary Material (lh_cVEL). The models simulate the pole dynamics accurately over the first *h* milliseconds (extrapolation horizon) and then assume constant pole velocity for the remaining time of the occlusion (900 *ms* − *h*). We investigated 61 values of the parameter *h*, corresponding to the 61 frames (Δt = 1/60s) from 100ms (*h* = −100*ms*) before to 900ms (*h* = *900ms*) after the pole occlusion. Notice, that the models for *h* = 0 *ms* and *h* = 900*ms* coincide with the previously specified models cVEL and PERF. The average prediction error of the models decreases monotonically with increasing *h* (Supplementary Material). Our interpretation of negative extrapolation horizons is that the respective subject solely relies on the constant velocity assumption to extrapolate the pole angle and additionally uses a deprecated estimate of the pole velocity (e.g. derived from 50ms before occlusion), instead of the actual velocity immediately before the occlusion.

We fitted the parameter *h* for each subject separately using the individual responses in each trial of block T4 (see Supplementary Material for details). Notice, that this procedure is different from fitting the model parameter based on the mean prediction errors. After having determined the best fitting model for each subject, we compared the groups also based on the parameter *h.* In analogy to the model class lh_cVEL, we also explored the class lh_cACC, which corresponds to assuming constant pole acceleration after first extrapolating the pole angle for *h* milliseconds accurately. However, this model class was not able to explain subjects’ behavior, because subjects made larger errors than the model could explain.

## Results

### Rating of balancing attempts is influenced by the balancing duration and mean absolute pole angle

During visual familiarization, every subject in the group VF rated overall 100 balancing attempts on a scale from one (very bad) to five (very good). In order to verify that subjects did not respond randomly, but paid attention to the balancing attempts and the cart-pole dynamics, we examined whether the rating of the subjects were meaningful. Two salient parameters for rating the balancing attempts are the balancing duration (BD) and the mean absolute pole angle (maPA).

First, a mixed-effects ANOVA was conducted, for subjects in the group VF, that examined the effect of rating for the two parameters, BD and maPA (**Figure 3**A, B). We found a significant effect of the factor rating for each of the parameters (p<0.001), suggesting that the rating separated each parameter into distinct classes. Post-hoc test revealed significant pairwise differences between all rating classes for BD (p<0.001, Holm corrected). For the maPA, all but one (3 vs 4) pairwise difference were significant (p<0.01, Holm corrected). Hence, subjects used the rating to classify the two performance measures, balancing duration and mean average pole angle, into ordered, significantly different, and thereby meaningful classes.

**Figure 3.**
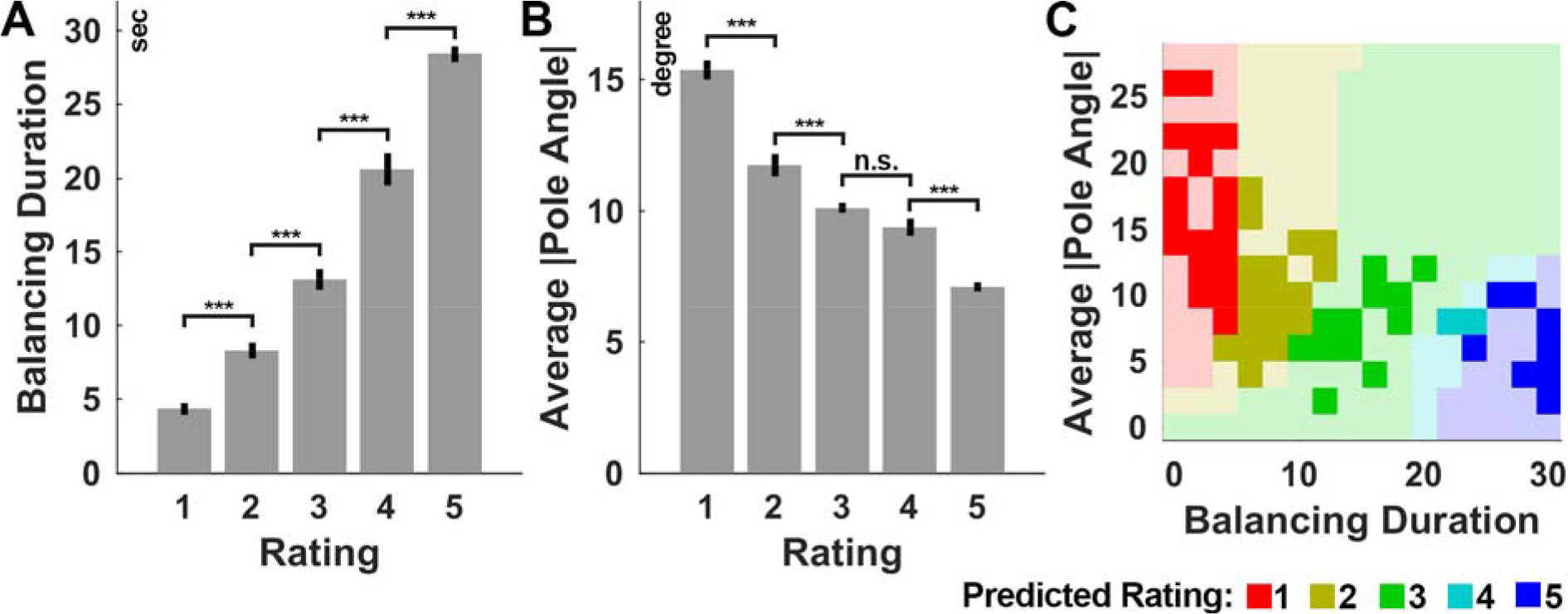
Analysis of subjects’ ratings during visual familiarization. (A) Balancing duration and (B) average absolute pole angle as function of subjects’ ratings. Data shown are the averages (bar) ± S.E.M. (C) Predicted rating of the fitted multinomial log-linear model as function of balancing duration and average absolute pole angle fitted to subjects’ responses. The model prediction was sampled uniformly in both dimensions for visualization. Opacity indicates whether training samples are close. Notice, that the rating (color) depends on both parameters, balancing duration and average absolute pole angle. *** p<0.001, n.s. p>0.1.

The influence of the two parameters (BD, maPA) on the rating was examined using a multinomial log-linear model (see Data Processing and Analysis). Both parameters and their interaction contributed significantly to the classification (all p<0.01). **Figure 3**C shows the predicted rating of the fitted model as function of the two parameters. Notice that the classification depends on both parameters. Inferring the mean absolute pole angle requires attentive observation throughout the whole trial. Thus, subjects in the group VF paid attention to the presentation of each balancing attempt and adhered to a non-trivial set of rules, based on the trial length and mean absolute pole angle, for classifying the balancing attempts into meaningful classes.

### Precision in extrapolation does not improve without feedback

For the pole extrapolation task, we first inspected the prediction error across blocks visually. Subjects of both groups show in average negative errors, which corresponds to a systematic underestimation of the pole movement (**Figure 4**A). This suggests that extrapolating the state-dependent acceleration of the pole is even for motor familiar subjects difficult. Furthermore, we noticed, that the variability between subjects in the group MF seems to be higher than for group VF. A possible reason could be a correlation between the extrapolation performance and the balancing proficiency, which varied between subjects although all motor familiar subjects were able to balance the cart-pole system (see Supplementary Material). However, we did not find any significant correlation between the average balancing duration in the motor control blocks and the prediction error in the subsequent extrapolation task block (for all blocks p>0.14, Spearman’s rank correlation). Furthermore, the between-subject variability was similar for both groups over the last blocks (F1-F5). Before comparing groups, we examined whether there is any improvement in extrapolating the pole angle when no feedback is provided (**Figure 4**A), which would suggest that subjects improve in pole extrapolation due to the rating (group VF) or control (group MF) task. Specifically, we conducted for each group a mixed-effects ANVOA that examined the effect of block on the prediction error over the blocks B and T1-T4. There was no significant effect of block in any of the groups (MF: p=0.87, VF: p=0.95). Hence, the ability to predict the pole angle does not improve without task specific feedback.

**Figure 4.**
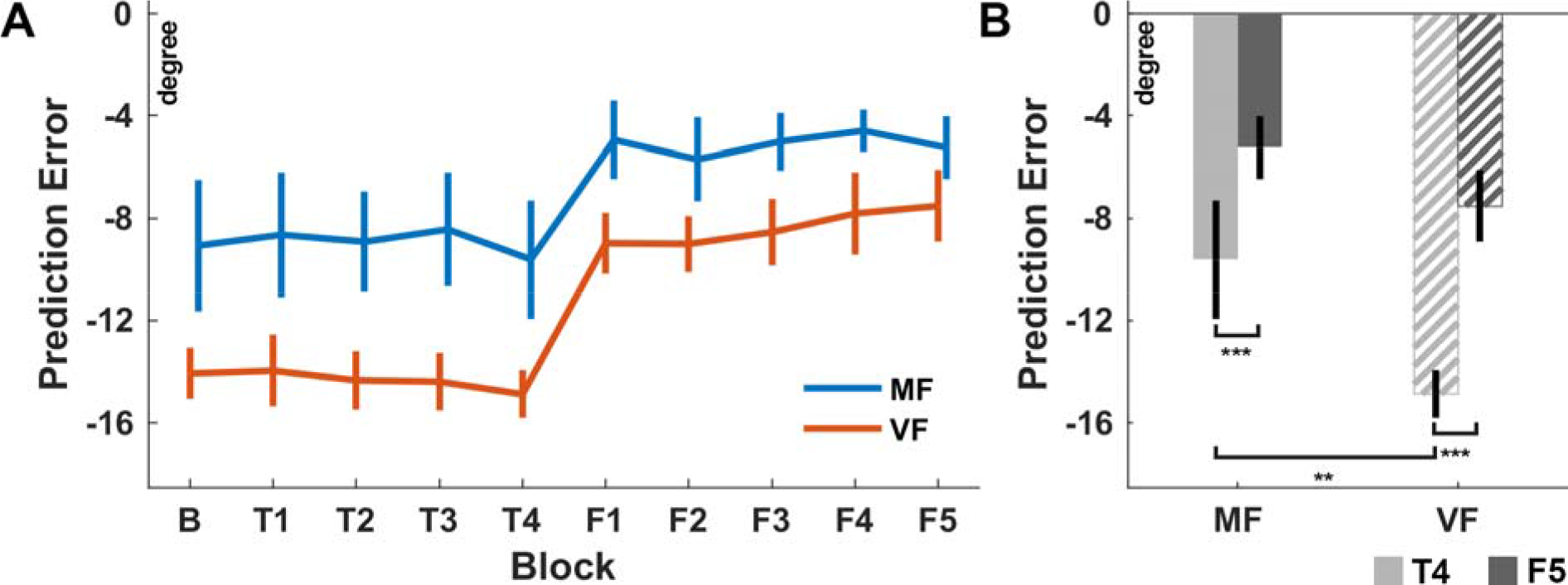
Average prediction error for both groups. (A) Average prediction error of the motor familiar (MF) and visually familiar (VF) group across all examined blocks. Negative errors correspond to an underestimation of the pole downwards movement. Notice that neither of the two groups improves significantly over the blocks B and T1-T4. Only during the blocks F1-F5, feedback is provided. (B) Prediction error of both groups in the blocks T4 (light gray) and F5 (dark gray). Within each group, subjects improve significantly due to feedback. There is a significant difference between the two groups before feedback was provided (T4). The prediction error is however not significantly different between groups after feedback was provided (F5). ** p<0.01, *** p<0.001. Error bars indicate ±1 S.E.M.

### Motor familiar subjects predict the pole angle more accurately

In order to test our hypothesis, that motor familiar subjects show a smaller prediction error than visual familiar subjects, we compared the prediction error between the groups just before any task specific feedback was provided (T4) and at the end of all feedback blocks (F5). To this end, a mixed-effect ANOVA was conducted that examined the effect of feedback (see Data Analysis) and group on the prediction error (**Figure 4**B). Both effects reached significance (group: p<0.02, feedback: p<0.001). The interaction did not reach significance (p=0.07). Post-hoc tests revealed a significant difference between groups before feedback was provided (p<0.01, Holm corrected, mean MF: −9.6°, VF: −14.9°) but not after (p=0.22, Holm corrected, mean MF: −5.2°, VF: −7.5°). Post-hoc comparison of the two blocks (T4: no feedback vs. F5: feedback) revealed a significant improvement within groups (both p<0.001). In summary, this analysis revealed a significant difference between the groups MF and VF before task specific feedback was provided. Thus, motor familiar subjects can predict the pole movement more accurately. Subjects in both groups are however able to utilize the feedback for improvement, resulting in statistically indistinguishable extrapolation performance in the last block (F5).

### Larger extrapolation horizon accounts for better extrapolation performance

We also investigated the average prediction error of different response models for extrapolating the pole angle during the occlusion (see Methods). Visual inspection reveals that, on average, both groups performed worse than a response model that accurately predicts the pole movement (PERF) or than assuming constant acceleration (cACC, **Figure 5**). For a more detailed analysis, we fitted subjects’ responses using our model class lh_cVEL, which extrapolates the state over the first *h* milliseconds (extrapolation horizon) accurately and then assumes constant pole velocity for the remaining time of the occlusion (see Methods and Supplementary Material). This model mimics the potentially limited capability to predict accurately over longer periods. In comparison to visually familiar subjects, behavior of motor familiar subjects was described by a significantly larger extrapolation horizon (parameter *h*, p=0.026, Wilcoxon rank sum test, median MF: 183.33ms, VF: 16.66ms, mean MF: 211.11ms, VF: 5.00ms). Thus, the subjects in the group MF seem to possess a more accurate, although not perfect, internal representation of the pole dynamics, which accounts better for the angle-depended acceleration of the pole.

**Figure 5.**
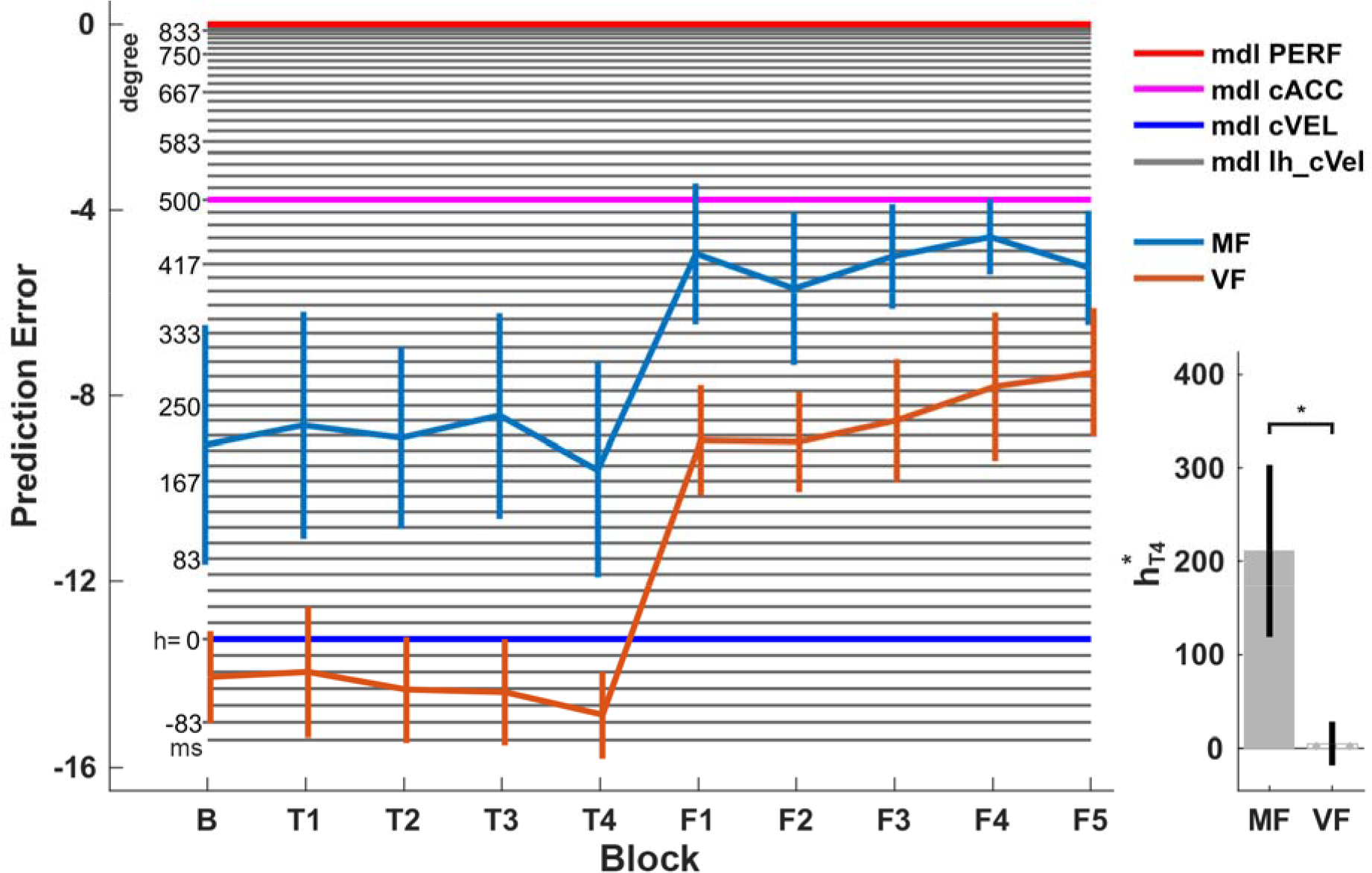
Average prediction errors for the investigated response models. In addition to the average prediction error for each model, the average prediction errors of subjects in the groups MF (light blue) and VF (orange) are shown. The three highlighted models (red: PERF, magenta: cACC, blue: cVEL) correspond to common assumptions in model-based extrapolation of motion (perfect model, constant acceleration and constant velocity). The average prediction error of each model in the class of partly heuristic forward models (lh_cVEL) is plotted as gray line, where h denotes the extrapolation horizon of the model. Notice that h = 0 and h = 900 coincide with the models cVEL and PERF. The model cACC is not in the class lh_cVEL. The parameters h^*^ of the models that fit subjects’ behaviors best in block T4 are significantly higher, and therefore closer to the perfect model (h = 900), for subjects in the group MF. Error bars indicate ±1 S.E.M.

## Discussion

We compared human’s ability to extrapolate the pole angle of the cart-pole system between subjects with and without motor expertise. Subjects without motor expertise were visually familiarized with the system in dedicated blocks during the experiment, while motor familiar subjects instead performed again the balancing task during this time as they have done previously during skill acquisition. During the extrapolation task, subjects observed balancing attempts and indicated for each attempt where they expected the pole to be after the pole was occluded for 900ms. Our analysis has revealed that neither of the two groups improves without receiving feedback about the true pole angle (**Figure 4**). Subjects with motor expertise were, however, even before receiving feedback significantly more precise in extrapolating the pole angle, although both groups received similar visual information during the familiarization blocks. We examined and explained the higher precision of motor familiar subjects in terms of exploiting a more accurate model of the pole dynamics, especially regarding the influence of the angle dependent gravitational acceleration of the pole and the ability to extrapolate this influence over a long duration accurately (extrapolation horizon, **Figure 5**). Subjects of both groups improved significantly in terms of minimizing the extrapolation error by using the feedback about the true pole angle in the last phase of the experiment (F1-F5). Finally, at the end of the experiment (F5), the extrapolation performance of the two groups was statistically indistinguishable.

La Scaleia et al. [10] argued that subjects incorporate prior experiences about gravity and air drag when intercepting ball trajectories in the form of model-based control. Since we are all used to the motion and behavior of objects during free fall, we already possess the knowledge that is necessary to catch falling objects. Similarly, we are used to human motion in the sense that we are able to interpret and extrapolate it [17]. It is important to note, that we usually acquire this ability and the corresponding knowledge implicitly and only rarely use it explicitly, meaning we use it for performing actions instead of expressing our expectations verbally. Nevertheless, subjects were able to discriminate plausible from implausible future body postures and to report their decisions [17]. Although the way subjects responded in the two studies differed substantially (actual catching vs. binary discrimination), the concept of internal simulation of the motion plays a central role for both studies in explaining subjects’ performances. These internal simulations of motion and the corresponding neural representations are arguably formed already during our childhood as result of throwing and catching balls, observing other people move as well as moving ourselves. During the training of specific tasks, such as playing basketball, these internal simulations are refined and specialized, which enables, for example, elite basketball players to outperform less experienced individuals in predicting the binary outcome of free shots [18]. Similarly, subjects with motor expertise were in our experiment more precise in predicting the pole angle than subjects without motor expertise. However, in comparison to Aglioti et al.’s experiment, the most important difference is that the cart-pole extrapolation task requires subjects to indicate the expected pole angle as precisely as possible instead of making only binary decisions (outcome of free shot). Thus, the underlying internal simulation has to fulfil substantially more restrictive accuracy requirements. In other words, the extrapolation of the pole angle has to be very precise in order to be able to discriminate between the diverse response options reliably. According to the internal simulation hypothesis [23], the precision of the extrapolation is determined by the accuracy of the internal forward model that represents the cart-pole system. We found that the responses of subjects with motor expertise are more similar to those of a perfect model than the responses of visual familiar subjects (larger extrapolation horizon, **Figure 5**). Thus, motor subjects seem to use a more accurate model for extrapolation. However, we also found that even motor familiar subjects are not able to extrapolate the motion without flaws. In fact, our model-based analysis suggests that subjects only extrapolate for about 211ms accurately (group MF in block T4, **Figure 5**) before switching to the constant velocity assumption. Similarly, Mehta and Schaal [15] reported that subjects show an average visuomotor delay of 269ms during virtual pole balancing and that trained subjects can additionally tolerate occlusions of 500-600ms duration without losing balance. Regarding the last point, it should however be noted that balancing the pole might be possible without being able to extrapolate the pole movement perfectly over the whole duration of the occlusion. In fact, since small mistakes during balancing can, to a certain degree, be corrected for later, subjects might acquiesce imperfections in order to reduce the effort for extrapolating more precisely. Nevertheless, when confronted with the error in the extrapolation task, subjects use the additional information to improve in this specific task.

Intriguingly, motor familiar subjects cannot only use the forward model, which they acquired during motor training, for performing actions predictively [14] but also to extrapolate and report the state of the system precisely, suggesting that information between action and perception is shared. Hecht et al. [24] have investigated the transfer of information and the relation between acting and perceiving in more detail. In their experiment subjects either had to rate (perceptual task) or perform (motor task) two subsequent movements that exhibited a certain relative timing. By permuting the sequence of tasks across groups, they investigated the transfer of knowledge from action to perception (action-perception transfer, APT) and vice versa (perception-action transfer, PAT). In comparison to a control group, both test groups were better in the second task, suggesting that information is transferred in both directions. Another interpretation is that for the perceptual and motor tasks the same representation is used and thereby the information is shared [25,26].

Recent research in the field of sensorimotor control addressed the topic of shared representations for action and perception from a slightly different perspective. Instead of acquiring a new motor skill, researchers investigated how the adaptation of motor behavior influences the perception. Sensorimotor adaptation is commonly explained by the adaptation of internal forward models that map actions to their sensory consequences. By minimizing the sensory prediction error (difference between predicted and actual outcome), predictions about the sensory consequences of actions are maintained accurate [27]. When the sensory consequences are artificially biased, for example due to a visuomotor rotation in virtual reality [27,28], the sensorimotor map is perturbed. Intriguingly, after having adapted to such perturbation as result of the minimization of the sensory prediction error, not only the subsequent motor execution is biased but also the perceived sensory consequences are biased in comparison to reality. Indeed, for classical visuomotor rotation paradigms, it has been shown that healthy subjects perceive their hand at a different position after sensorimotor adaptation and that the perceived hand position corresponds to the visuomotor rotation [29]. These studies suggest that extrapolation mechanisms for motor control and perception are not only tightly related but might also share the same internal forward models. Regarding the neural correlates of this behavioral finding, Synofzik et al. [29] have shown that not only the adaptation of motor behavior [16] but that also the shift in perception depends on the integrity of the cerebellum. Especially the posterior-lateral cerebellum seems to be important not only for the planning of motor actions [16,30] but also for the perception of action [31,32] and sensory prediction [5–9,33,34]. It seems therefore plausible that the cerebellum implements mechanisms, which are equally used for perception and action. Whether the cerebellum and other brain areas, such as the parietal lobe [35–39], are involved in the cart-pole balancing and extrapolation tasks, and whether there are different neural correlates depending on the preceding type of training (visual vs. motor), is an interesting direction for future research.

Finally, we would like to address a limitation of this study, which is that motor familiar subjects interacted overall for a longer duration with the cart-pole system due to the prior skill acquisition. However, although we were not able to perform a statistical comparison, the control experiment suggests that the duration of visual exposure does not influence prediction accuracy in this specific task. Furthermore, the goal of this study was not only to reveal a general behavioral difference between visual or motor familiar subjects, as has similarly been reported before [18,24], but also to provide a model-driven analysis that characterizes and explains individual prediction performances.

## Conclusions and Outlook

In summary, we have shown that motor training does not only improve motor control but also task-relevant perceptual abilities. Being able to extrapolate the state of the cart-pole system into the future is an important ability for easeful balancing performance. Correspondingly, subjects who had previously learned to control the cart-pole system were more precise in extrapolating the pole movement. Our results suggest that motor training yields an accurate, although not perfect, internal forward model of the controlled dynamics, which can be used for both, controlling and accurately extrapolating the dynamic behavior of the cart-pole system. Similar extrapolation accuracy can, however, be achieved without motor training by minimizing the prediction error when task-specific feedback is provided. Since our model explains the difference in extrapolation performance based on the duration of accurate simulation, a future study could tested this prediction explicitly by varying the duration of the occlusion. Another interesting question for future investigations is whether the ability to extrapolate the pole movement more precisely improves the cart-pole balancing performance in the case of motor familiar subjects or facilitates the acquisition of the cart-pole balancing skill for visual familiar subjects (perception-action transfer, similar to Hecht et al. [24]). Furthermore, comparison of the neural correlates corresponding to cart-pole balancing and extrapolating the pole movement might foster the understanding of knowledge representations in the brain and their acquisition.

## Acknowledgements

The authors would like to thank Andrea Christensen for the valuable comments and suggestions to improve the manuscript. NL would like to thank the German National Academic Foundation for granting a doctoral student fellowship. Financial support for this study has been received from the German Research Foundation (DFG GZ: KA 1258/15-1), European Union Seventh Framework Programme (CogIMon H2020 ICT-644727) and the Human Frontiers Science Program (HFSP RGP0036/2016). The authors declare no competing financial interests.

## References

1. Franklin DW, Wolpert DM. Computational Mechanisms of Sensorimotor Control. Neuron. 2011; 72: 425–442. doi: 10.1016/j.neuron.2011.10.006.

2. Wolpert DM, Diedrichsen J, Flanagan JR. Principles of sensorimotor learning. Nature Reviews Neuroscience. 2011; 12: 739–751. doi: 10.1038/nrn3112.

3. Yarrow K,Brown P, Krakauer JW. Inside the brain of an elite athlete: the neural processes that support high achievement in sports. Nature Reviews Neuroscience. 2009; 10: 585–596. doi: 10.1038/nrn2672.

4. Zago M,McIntyre J,Senot P, Lacquaniti F. Visuo-motor coordination and internal models for object interception. Experimental Brain Research. 2009; 192: 571–604. doi: 10.1007/s00221-008-1691-3.

5. Engel A,Burke M,Fiehler K,Bien S, Rosler F. Motor learning affects visual movement perception. Eur J Neurosci. 2008; 27: 2294–2302. doi: 10.1111/j.1460-9568.2008.06200.x.

6. Ivry RB,Spener RM,Zelaznik HN, Diedrichsen J. The Cerebellum and Event Timing. Annals NY Acad Sci. 2002; 978: 302–317. doi: 10.1111/j.1749-6632.2002.tb07576.x.

7. Schubotz RI. Prediction of external events with our motor system: towards a new framework. Trends Cogn Sci. 2007; 11: 211–218. doi: 10.1016/j.tics.2007.02.006.

8. O’Reilly JX,Mesulam MM, Nobre AC. The cerebellum predicts the timing of perceptual events. J Neurosci. 2008; 28: 2252–2260. doi: 10.1523/JNEUROSCI.2742-07.2008.

9. Roth MJ, Synofzik M, Lindner A. The Cerebellum Optimizes Perceptual Predictions about External Sensory Events. Current Biology. 2013; 23: 930–935. doi: 10.1016/j.cub.2013.04.027.

10. La Scaleia B,Zago M, Lacquaniti F. Hand interception of occluded motion in humans: a test of model-based vs. on-line control. Journal of Neurophysiology. 2015; 114: 1577–1592. doi: 10.1152/jn.00475.2015.

11. Dingwell JB,Mah CD, Mussa-Ivaldi FA. Manipulating Objects With Internal Degrees of Freedom: Evidence for Model-Based Control. Journal of Neurophysiology. 2002; 88: 222–235.

12. Nagengast AJ,Braun DA, Wolpert DM. Optimal Control Predicts Human Performance on Objects with Internal Degrees of Freedom. PLoS Comput Biol. 2009; 5. doi: 10.1371/journal.pcbi.1000419.

13. Hasson CJ, Shen T, Sternad D. Energy margins in dynamic object manipulation. Journal of Neurophysiology. 2012; 108: 1349–1365. doi: 10.1152/jn.00019.2012.

14. Ludolph N, Giese MA, Ilg W. Interacting Learning Processes During Skill Acquisition. Learning To Control With Gradually Changing System Dynamics. bioRxiv. 2017. doi: 10.1101/135095.

15. Mehta B, Schaal S. Forward Models in Visuomotor Control. Journal of Neurophysiology. 2002; 88: 942–953.

16. Donchin O,Rabe K,Diedrichsen J,Lally N,Schoch B, Gizewski ER, et al. Cerebellar regions involved in adaptation to force field and visuomotor perturbation. Journal of Neurophysiology. 2011; 107: 134–147. doi: 10.1152/jn.00007.2011.

17. Graf M,Reitzner B,Corves C,Casile A,Giese M, Prinz W. Predicting point-light actions in realtime. Neuroimage. 2007; 36 Suppl 2: T22–32. doi: 10.1016/j.neuroimage.2007.03.017.

18. Aglioti SM, Cesari P,Romani M, Urgesi C. Action anticipation and motor resonance in elite basketball players. Nat. Neurosci. 2008; 11: 1109–1116. doi: 10.1038/nn.2182.

19. Brainard DH. The Psychophysics Toolbox. Spatial Vision. 1997; 10: 433–436. doi: 10.1163/156856897X00357.

20. Pelli DG. The VideoToolbox software for visual psychophysics: transforming numbers into movies. Spatial Vision. 1997; 10: 437–442. doi: 10.1163/156856897X00366.

21. Kleiner M, Brainard DH, Pelli DG. What’s new in Psychtoolbox-3. European Conference on Visual Perception; 2007.

22. Barto AG,Sutton RS, Anderson CW. Neuronlike adaptive elements that can solve difficult learning control problems. IEEE Transactions on Systems, Man, and Cybernetics. 1983; 13: 834–846. doi: 10.1109/TSMC.1983.6313077.

23. Grush R. The emulation theory of representation: Motor control, imagery, and perception. Behav. Brain Sci. 2004; 27. doi: 10.1017/S0140525X04000093.

24. Hecht H,Vogt S, Prinz W. Motor learning enhances perceptual judgment. A case for action-perception transfer. Psychological Research. 2001; 65: 3–14. doi: 10.1007/s004260000043.

25. Prinz W. Perception and Action Planning. European Journal of Cognitive Psychology. 1997; 9: 129–154. doi: 10.1080/713752551.

26. Casile A, Giese MA. Nonvisual motor training influences biological motion perception. Curr. Biol. 2006; 16: 69–74. doi: 10.1016/j.cub.2005.10.071.

27. Shadmehr R, Smith MA, Krakauer JW. Error Correction, Sensory Prediction, and Adaptation in Motor Control. Annu. Rev. Neurosci. 2010; 33: 89–108. doi: 10.1146/annurev-neuro-060909- 153135.

28. Prablanc C,Tzavaras A, Jeannerod M. Adaptation of hand tracking to rotated visual coordinates. Perception & Psychophysics. 1975; 17: 325–328. doi: 10.3758/BF03203218.

29. Synofzik M,Lindner A, Thier P. The cerebellum updates predictions about the visual consequences of one’s behavior. Curr. Biol. 2008; 18: 814–818. doi: 10.1016/j.cub.2008.04.071.

30. Higuchi S,Imamizu H, Kawato M. Cerebellar Activity Evoked By Common Tool-Use Execution And Imagery Tasks: An Fmri Study. Cortex. 2007; 43: 350–358. doi: 10.1016/S0010- 9452(08)70460-X.

31. Calvo-Merino B, Grèzes J, Glaser DE, Passingham RE, Haggard P. Seeing or doing? Influence of visual and motor familiarity in action observation. Curr. Biol. 2006; 16: 1905–1910. doi: 10.1016/j.cub.2006.07.065.

32. Christensen A, Giese MA, Sultan F, Mueller OM,Goericke SL, Ilg W, et al. An Intact Action-Perception Coupling Depends on the Integrity of the Cerebellum. Journal of Neuroscience. 2014; 34: 6707–6716. doi: 10.1523/JNEUROSCI.3276- 13.2014.

33. Blakemore S-J, Frith CD, Wolpert DM. The cerebellum is involved in predicting the sensory consequences of action. Neuroreport. 2001; 12: 1879–1884. doi: 10.1097/00001756-200107030- 00023.

34. Deluca C,Golzar A,Santandrea E,Lo Gerfo E,Estocinova J,Moretto G, et al. The cerebellum and visual perceptual learning: evidence from a motion extrapolation task. Cortex. 2014; 58: 52–71. doi: 10.1016/j.cortex.2014.04.017.

35. Wolpert DM,Goodbody SJ, Husain M. Maintaining internal representations. The role of the human superior parietal lobe. Nat Neurosci. 1998; 1: 529–533. doi: 10.1038/2245.

36. Desmurget M, Grafton S. Forward modeling allows feedback control for fast reaching movements. Trends Cogn Sci. 2000; 4: 423–431. doi: 10.1016/S1364-6613(00)01537-0.

37. Vahdat S,Darainy M, Milner TE, Ostry DJ. Functionally Specific Changes in Resting-State Sensorimotor Networks after Motor Learning. Journal of Neuroscience. 2011; 31: 16907–16915. doi: 10.1523/JNEUROSCI.2737-11.2011.

38. Vahdat S,Darainy M, Ostry DJ. Structure of plasticity in human sensory and motor networks due to perceptual learning. The Journal of Neuroscience. 2014; 34: 2451–2463. doi: 10.1523/JNEUROSCI.4291-13. 2014.

39. Blakemore S-J, Sirigu A. Action prediction in the cerebellum and in the parietal lobe. Experimental Brain Research. 2003; 153: 239–245.doi: 10.1007/s00221-003-1597-z.

